# Reviving collapsed networks from a single species: the importance of trait variation and network architecture

**DOI:** 10.1101/2023.09.30.560140

**Authors:** Gaurav Baruah, Meike Wittmann

## Abstract

Mutualistic ecological networks can suddenly transition to undesirable states, due to small changes in environmental conditions. Recovering from such a collapse can be difficult as reversing the original environmental conditions may be infeasible. Additionally, such networks can also exhibit hysteresis, implying that ecological networks may not recover. Here, using a dynamical eco-evolutionary framework, we try to resurrect mutualistic networks from an undesirable low-functional collapse state to a high-functioning state. We found that restoring the original environmental conditions rarely aided in recovering the original network due to the presence of hysteresis. By combining concepts from signal propagation theory and eco-evolutionary dynamical modeling, we show that network resurrection could be readily achieved by perturbing a single species that controls the response of the dynamical networks. We show that during the resurrection of collapsed networks, the historical network architecture, levels of trait variation, and eco-evolutionary dynamics could aid in the revival of the network even in undesirable environmental conditions. Our study argues that focus should be applied to a few species whose dynamics one could steer to resurrect the entire network from a collapsed state.

## 1 Introduction

When environmental conditions cross certain thresholds, complex ecological communities can exhibit abrupt transitions from a stable desirable community state to an undesirable state where the loss of ecosystem functioning and services occurs (1, 2). Examples are shifts of shallow lakes from clear to turbid state (3), the collapse of fisheries (4), or the collapse of vegetation leading to a deserted state (5). In addition to this, mutualistic networks governed by positive feedback loops are critically vulnerable to changes in environmental conditions (6, 7, 8). Loss of such complex networks or shifts to undesirable states would lead to a loss in ecosystem functions and services provided by them. Thus we need to understand the occurrence of such tipping points with an aim to reverse such undesirable effects.

Phenomenological statistical tools such as early warning signals have been suggested to help in forecasting such unwanted transitions (9, 10, 11, 12). However, there are many drawbacks associated with using such tools as most often these signals imperfectly forecast transition (13, 14). In case when such an abrupt transition occurs, a solution to recover such a system would be to restore the original environmental conditions where possible. However, this solution might not always work (15). More often than not we do not have information or understanding of the system and its properties at the brink of the collapse. Furthermore, such ecological systems could potentially exhibit hysteresis whereby the ecological system could occupy both functional and dysfunctional states for the same environmental conditions, making recovery to a fully functional state a difficult ecological problem (15, 16). It is thus crucial to understand how a dynamical ecological network behaves at the brink of a collapse in order to steer it back to its original functionality once such a network collapses. Previous studies have tried to understand how a dynamical system behaves in the vicinity of a transition (17). However, recovering an ecological network from an undesirable state is another problem that has somewhat remained overlooked. To fill this gap, one solution would be to understand what set of structural network properties (17, 18) or eco-evolutionary parameters could be pertinent in the reconstruction of a collapsed ecological network. Once this is known, it would then be easier to comprehend the parameter spaces at which an entire network could be resurrected and the steps that would be needed to ensure that.

Two important metrics quantifying the structure of an ecological network are connectance and nestedness (19, 20). Connectance quantifies how well species in an ecological network interact with other species, and nestedness quantifies species interactions of specialists with only species that interact with both generalists and specialists, i.e., how hierarchical the community is in terms of species interactions. These network properties govern how mutualistic networks respond to changes in environmental conditions such as species loss or decrease in average mutualism strength (21, 22, 17, 23). Previous studies have suggested that such network properties are pertinent to maintaining biodiversity and can impact stability, feasibility, and critical transitions (24, 21, 17). While such network features are known to be important for maintaining the stability of mutualistic networks, their role in the resurrection of collapsed networks remains under-explored. It is possible that while resurrecting collapsed low-functional networks, some networks never recover to their original functional state, which could be due to the arrangement of species interactions.

In a rapidly changing world, the maintenance of biodiversity is a key target (25). However, as the human footprint on our planet continues to grow, leading to widespread biodiversity loss and accelerating extinction rates (26, 27), forecasting biodiversity collapse should not be the only goal. We thus propose to investigate the resurrection of low-functional collapsed mutualistic networks by utilizing a phenomenon of propagation of perturbation in such ecological networks. For instance, a previous study (18) evaluated the spread of a single perturbation on a node over entire networks, finding that such signal spread can be dictated by structural factors of the networks. Since mutualistic ecological networks consist of species interacting in a certain arrangement, and because of positive feedback between species in such networks, perturbation propagation could possibly be used to resurrect networks that have transitioned to an undesirable state. This is akin to a local environmental disturbance that can spread through an entire ecological network and cause local extinctions, or a perturbation signal that spreads through components of various interaction networks (18).

Species embedded in complex ecological networks interact with each other through their phenotypic traits (28, 29). For instance, in mutualistic networks, such as a plant-pollinator network, successful and beneficial mutualism could be possible through matching phenotypes such as proboscis of pollinators and corolla lengths of plants (30, 31). If species possess similar matching phenotypes, successful interaction could lead to fitness benefits for both interacting species. In such a case, these interacting species possess intraspecific variation in such phenotypes that become crucial in maintaining interacting intimacy with multiple interacting species (32). Such individual variation, although demonstrated to be widespread in empirical studies, has largely been ignored in understanding its dynamical consequences on stability and resilience (but see 33, 12, 23). Recently, individual trait variation in species interaction networks has been suggested to have a significant impact on the occurrence of tipping points and abrupt collapses (23). Incorporating such trait variation in classical phenomenological models could help us understand how networks could respond to environmental perturbation and whether the presence of trait variation could aid in network resurrection when such networks collapse.

In this study, using a dynamical eco-evolutionary framework, we try to revive mutualistic networks from an undesirable alternative stable state to a high-functioning stable state at unfavorable conditions. We address how the original architecture of such networks, the arrangement of ecological interactions, and the presence of trait variation could aid in the resurrection. We found that restoring the original environmental conditions rarely aided in recovering the original network due to the presence of hysteresis. By combining frameworks from signal propagation theory in networks, and eco-evolutionary dynamical modeling, we show that network revival could be readily achieved by perturbing a single species that controls the response of such dynamical networks. We show that during the resurrection of such collapsed networks, the historical network architecture, levels of trait variation, and eco-evolutionary dynamics could aid in the revival of the network even at undesirable parameter spaces. Our study indicates that restoring original environmental conditions will rarely lead to the recovery of large mutualistic communities, but focus should be instead applied to a few species whose dynamics could steer the entire network to resurrection. In ecological practice, this would relate to increasing the survival of a particular species by some amount, or by adding individuals, or by controlling the density of a species constantly for a period of time.

## 2 Results

We used 115 mutualistic plant-pollinator networks from the web-of-life database (www.web-of-life.es). The networks comprised a total of 7159 species, with a maximum number of 167 species in a network and a minimum of 8 species. Nestedness (NODF) ranged from as low as 0 to as high as 0.84, while network connectance ranged from a low of 0.035 to a maximum of 0.58. Using the modeling framework detailed in section 4, we first evaluated the collapse regime of the mutualistic networks. That is, we identified for which values of the average mutualistic strength, *γ*_0_, the network collapses, if *γ*_0_ is set to this value. Next, we investigated whether restoring the original conditions could help recover the mutualistic networks, i.e., if these networks exhibited hysteresis. Finally, we evaluated through our modeling framework, whether positively perturbing one focal species in a network could help revive collapsed networks even in unfavorable environmental conditions.

### 2.1 Collapse of networks and hysteresis

Among 115 mutualistic networks analyzed, the collapse regime, i.e., the parameter region at which the networks transitioned to the collapse state (i.e., less than 90 percent of its original richness) was *γ*_0_ *≤* 1.5 (Fig. 1D-E). Thus,*γ*_0_ *≤* 1.5 was considered unfavorable environmental conditions. Out of 115 networks analyzed, 94 percent of the networks exhibited strong hysteresis i.e., the networks never recovered even at high values of *γ*_0_ = 4.85, and recovery never occurred despite reverting to the original environmental conditions (Fig 1B, 1C, Fig. S1).

**Figure 1:**
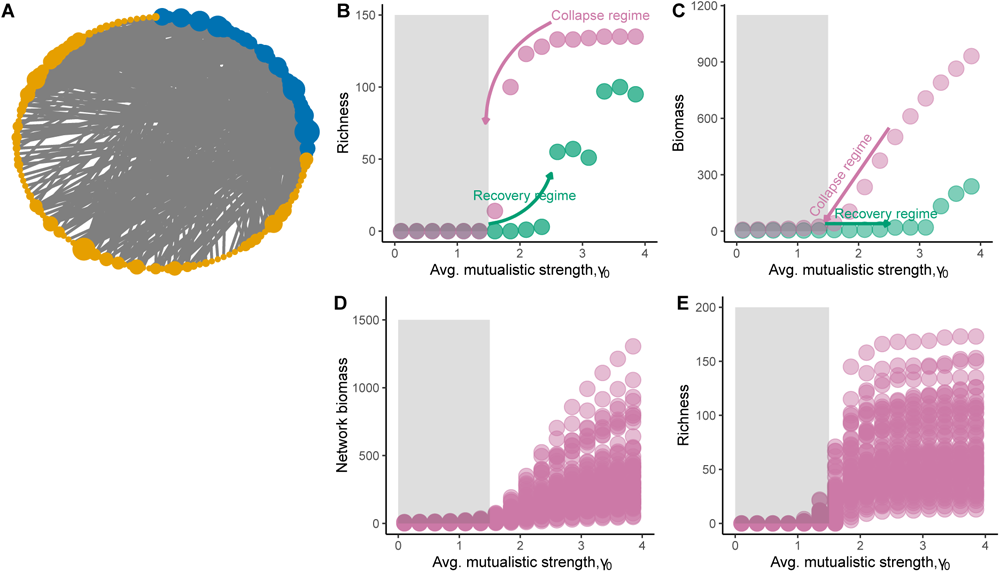
Hysteresis and collapse regime in mutualistic networks. (A) An example mutualistic plantpollinator network of 142 species. The mutualistic network shown here has nestedness of 0.19 and connectance of 0.09. The size of the nodes indicates the number of unique intergroup interactions i.e., the larger the node size is, the higher is the number of interactions. (B) As the transition parameter, which is the average strength of mutualism, *γ*_0_, decreases the mutualistic network transitions from a high-functioning stable state – high species richness in B and high biomass in C – to a low functioning undesirable state – low richness in B and low biomass depicted by dark pink points. This is the collapse regime, i.e., the mutualistic network completely collapses in the region of, *γ*_0_ = [0, 1.5]. However, as the environmental conditions are reversed gradually, i.e., mutualistic strength is increased slowly (shown by the dark green points), network recovery does not materialise as the system stays at the undesirable low functioning state (dark green points) and thus exhibits hysteresis. D-E) For all 115 mutualistic plant-pollinator networks, the collapse regime (dark pink points) occurs around *γ*_0_ *∈* [0, 1.5]. The grey-shaded region in B, C, D, E represents this collapse regime and the regime where we directed all further analysis.

### 2.2 Network resurrection from a single species: role of trait variation and evolutionary dynamics

When choosing the species with the highest degree for positive perturbation, mutualistic networks could be revived even at a very low mutualistic strength of *γ*_0_ = 1.15 (Fig. 2, Fig. 3F-G). Figure 2 shows an example resurrecting dynamics of two mutualistic networks: one with 34 species and another with 49 species. At very low *γ*_0_ = 1.15, mutualistic networks transition to the collapse state and without any intervention remain at the undesirable collapse state (Fig. 1D-E, Fig. 2B, 2E). With 0.5 forcing strength on the species with the highest degree in both networks for a duration of time (*T* = 500), both the networks easily recovered back to high functionality (Fig. 2C, 2F) at low *γ*_0_ = 1.15. For another large network of 112 species, high trait variation can aid in resurrecting collapsed networks from a single species perturbation (Fig. 3A-D). With low trait variation, the spread of perturbation from the species with the highest degree played no significant role in resurrecting collapsed networks i.e., the spread of perturbation was not strong enough to positively impact densities of interacting species (Fig. 3E-H). The presence of evolutionary dynamics also impacted the resurrection of collapsed networks from a single species (supplementary Fig. S3-S5). When evolution was turned off, i.e., heritability was fixed at *h*^2^ = 0 for all species, network resurrection from perturbing the most generalist species irrespective of high or low trait variation, failed. However, this changes the moment evolutionary dynamics come at play, i.e., when heritable variation is no longer zero but *h*^2^ = 0.4 for all species, and trait variation was high (supplementary Fig. S3-S4). Although, phenotypic trait change plays an important role, even when heritability was zero, i.e., in the absence of evolutionary dynamics, network revival from a single species is a possibility but in a restricted parameter space of the collapse regime (supplementary Fig. S5).

**Figure 2:**
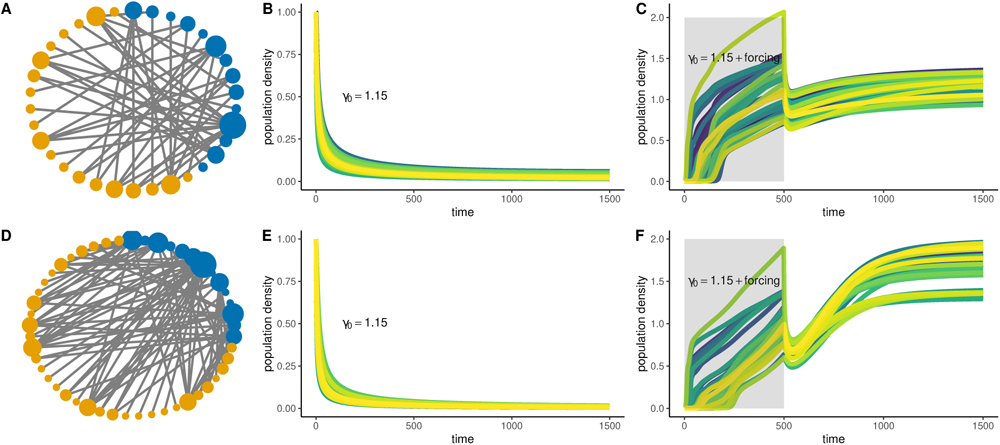
Network resurrection from perturbing a single species in two contrasting networks of different architecture. A) A mutualistic plant-pollinator network with 34 species with nestedness of 0.27 and connectance of 0.15 and D) a network with 49 species with nestedness of 0.32 and connectance of 0.17. (B, E) Both networks transition to a low-functioning undesirable state at low average mutualistic strength of *γ*_0_ = 1.15. In B and E, initial species densities were fixed at 1, and trait variation for all species for both the networks was fixed at *σ_i_* = 0.02. Initial mean trait values *u_i_* were based on table 1 values. (C, F) Perturbing the species with the highest degree in each of the two networks with a forcing strength of 50 percent at the same average mutualistic strength of *γ*_0_ = 1.15, for a duration of 500 time points leads to the resurrection of the two networks i.e., all *N_i_ >* 0.5. Here for C and F, we considered initial starting species densities to be *N_i_ <* 0.005 and trait variation for all species to be at a moderate level of *σ_i_* = 0.02.

**Table 1:** Parameters and their values with the description used for revival of collapsed networks.

| Symbol | Description |
| --- | --- |
| $b$ | Intrinsic growth rate which was fixed at 0 for all species indicating obligate mutualism. |
| $\sigma_i$ | Trait standard deviation of species $i$; fixed at 0.005 for low trait variation and 0.02 for high trait variation. |
| $h_i^2$ | heritable variation of species $i$'s trait, fixed at 0.4. |
| $\gamma_0$ | Average mutualistic strength among species that was in the range of 0.5 to 1.5 which was in the collapse range (see fig. 1) |
| $d_i$ | Degree of species $i$ in a network |
| $H$ | Handling time of species which was fixed at 0.25 for all species |
| $A_{ik}$ | Adjacency matrix entry for the interaction between animal species $i$ and plant species $k$ that is either 0 or 1. |
| $\alpha_{ij}$ | Competition matrix entry where intraspecific competition $\alpha_{ii} > \alpha_{ij}$. $\alpha_{ii}$ was fixed at 1 and $\alpha_{ij}$ was drawn from uniform distribution of $U[0.0001, 0.001]$ |
| $\omega$ | Width of the pairwise mutualistic Gaussian kernel fixed at 0.35. |
| $u_i$ | Mean phenotypic trait values of species. The initial starting values were randomly sampled from $U[-0.5, 0.5]$. The final values used to determine whether networks could be revived were used when networks were at quasi-positive equilibrium. |
| $N_i^{(A)}, N_i^{(P)}$ | Density of species (plants and pollinators) from the collapsed state and drawn from a random uniform distribution on $U[0, 0.005]$ mimicking a state where all species had density either 0 or very low. |
| $\nu(T)$ | Forcing strength of a species for a duration of time given by $T$. Forcing strength varied from 0 to 1, 0.5 for figure 2, and varied for figure 3. Duration $T$ also varied from 100 to 500. |

**Figure 3:**
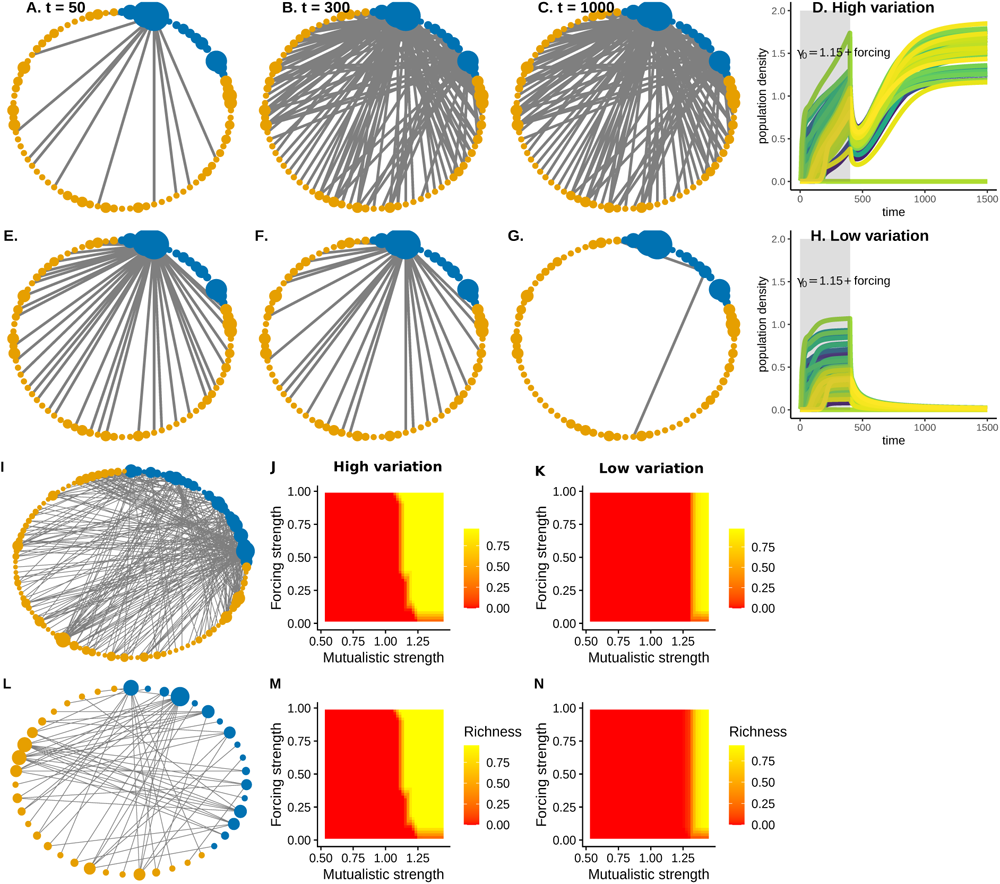
Role of trait variation for the revival of a network from a single species. (A-D) High trait variation, i.e., *σ_i_* = 0.02: A network with 112 species is being reignited from an undesirable stable state i.e., all species had *N_i_ <* 0.0005 and *γ*_0_ = 1.15. For resurrecting the collapsed network, a single species with the highest number of interactions was positively perturbed repeatedly for a duration of 400 time points at a forcing strength of 0.5. (A-C) Time evolution of how the recovery of the network takes place (D) Population dynamics and successful resurrection over time as the species with the highest degree is perturbed. (E-H) Low trait variation case, i.e.,*σ_i_*= 0.005. (E) Initially, at *t* = 50 due to perturbation that spread through the network, species recovered their inter-group connections. However, this recovery was temporary as the network fails to recover as shown in (G) *t* = 1000 and in (H). (I-K) The parameter space for successful resurrection for another network with 112 species differed for the case when species had high trait variation versus when species exhibited low trait variation. For the network with 112 species only at high forcing strength that was greater than 0.25 resurrection of the network was possible at low average mutualistic strength, *γ*_0_ *≤* 1.25. (K) For low trait variation, only at high *γ*_0_ *≥* 1.25, the resurrection was possible. (L-N) For a slightly smaller network with 46 species, we observed similar results. Successful resurrection was possible for high trait variation at low *γ*_0_ *<* 1.2 provided there was a slightly high forcing strength.

### 2.3 Network resurrection from a single species: role of network architecture

As network structure varied, resurrection from a collapsed state to a functional state also varied (Fig. 4). Recovery species richness (i.e., all *N* ^(^*^A,P^* ^)^ *>* 0.5 density) after perturbing the species with the highest degree was higher when nestedness and connectance of networks were high. Networks that are either small in size, or networks that are high in nestedness or connectance recover to their functional form with only 0.5 forcing strength (*ν*) applied to the species with the highest degree (Fig. 4A-C). This was observed more so when species exhibited some amount of trait variation (Fig 4). Furthermore, mean species density achieved in each of the 115 networks at low levels of *γ*_0_ after the perturbation of the most generalist species was stopped was substantially higher when species exhibited high trait variation (Fig. 5B). In addition, the proportion of species in each of the 115 networks that achieved a density greater than 0.5 after species-specific perturbation was stopped after a certain duration *T* = 500, was higher when species had high trait variance (Fig. 5C). This result, however, was dependent on the architecture of the networks. In other words, at low levels of mutualistic strength, *γ*_0_ *<* 1.2, and high levels of trait variation, the proportion of species that achieved higher density was positively correlated with how nested the networks were (Fig. 5A). In additional analysis, and in conjunction with perturbation being applied to the species with the highest degree, we perturb an additional parameter conducive to the spread of the perturbation, which was the intrinsic growth rate of the species. However, all other *S −* 1 species’ growth rates remain as they are. As shown in the supplementary material with three contrasting networks, perturbing the rate of growth of the most generalist species alongside a constant perturbation of 50 percent for a period of 500 time points does not always lead to successful resurrection. Controlling the rate of growth had minimal impacts on resurrection for larger networks (supplementary Fig. S6). In addition, adding a certain density of a species for a certain duration could also be beneficial for reviving collapsed networks (supplementary Fig. S7-S8).

**Figure 4:**
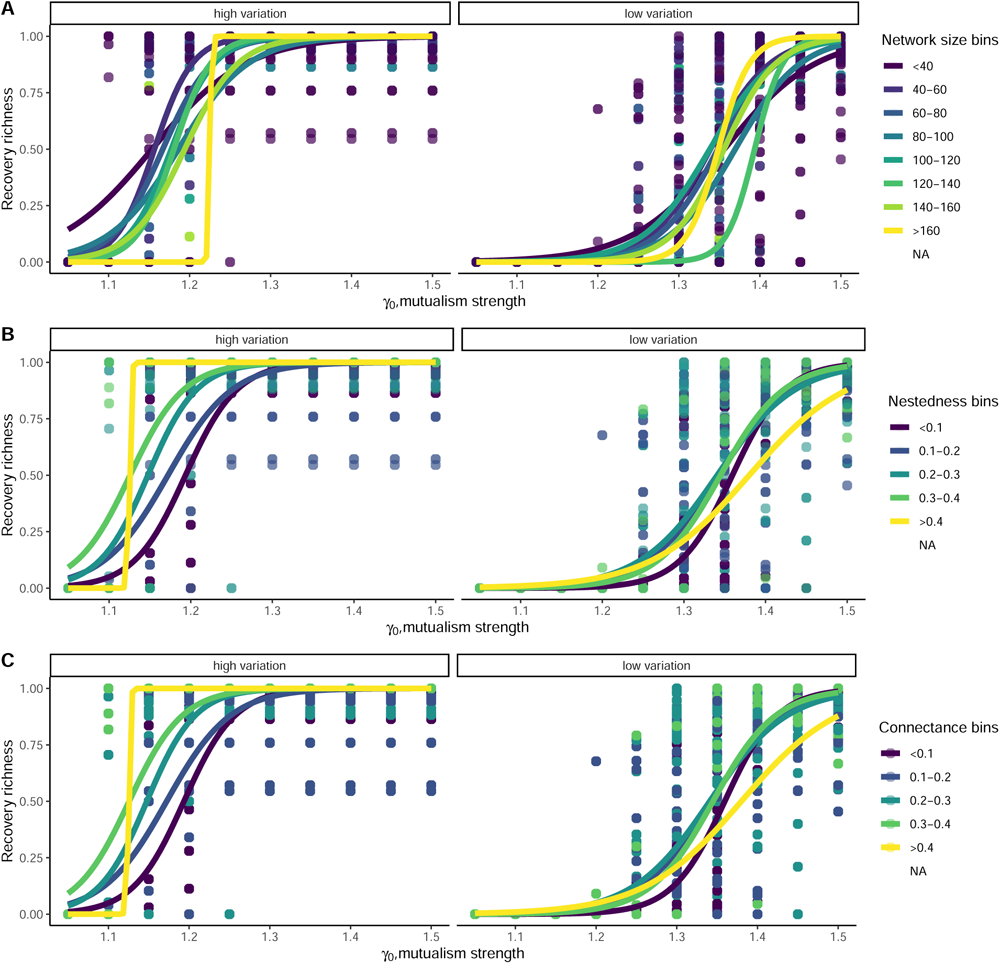
Network recovery from perturbing a single species was impacted negatively by network size (A), positively by nestedness (B), and connectance (C), particularly when species had higher trait variation. In each of these networks, only the species with the highest degree, was positively perturbed from a very low density, *N_i_ <* 0.005, for a duration of 500 time points with a forcing strength of 0.5, while the rest of the species remained unperturbed. Shown here are data from 115 networks. *σ_i_* was fixed at 0.02 for the high trait variation case and 0.005 for the low trait variation case respectively. Initial mean trait values were sampled according to parameter values given in table 1.

**Figure 5:**
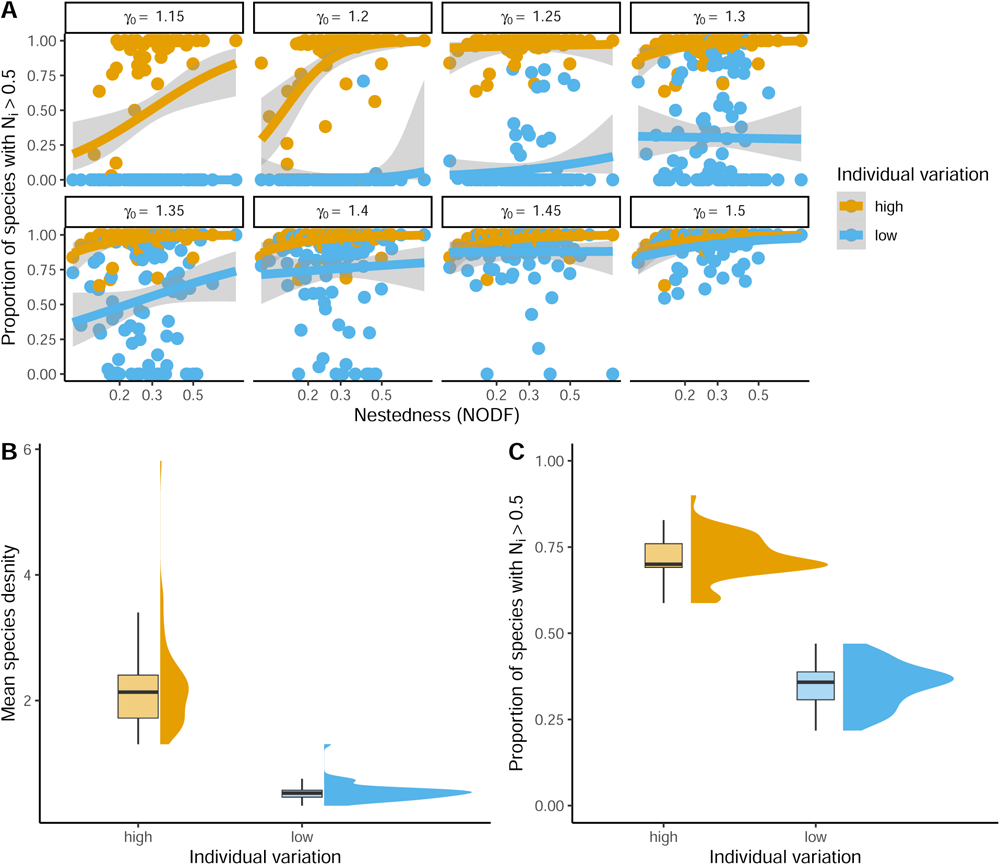
Nestedness (NODF) and individual variation impacted network recovery and species responses. (A) As nestedness increased, the proportion of species in each network with a density higher than 0.5, after the focal species perturbation was stopped, increased more so when species had high trait variation in contrast to when species had low trait variation, which was particularly evident at low *γ*_0_ *<* 1.35. The proportion of species in each network reaching higher density was significantly lower for low trait variation. In all these cases, only one species was forced at a forcing strength of 0.5 and for a duration of 500 time points. Each point in (A) represents a mutualistic network, a total of 115 networks are shown in this figure. The thick lines represent a generalized linear model fit with quasibinomial error distribution and the shaded grey region represents standard error. (B) Mean species density at equilibrium, after a forcing duration of 500 time points, was significantly higher when networks had high trait variation in comparison to networks that had low variation. (C) in the case of high trait variation, for all *γ*_0_ *>* 1, 76 percent of species in each network had a density higher than 0.5 i.e., *N_i_ >* 0.5 at equilibrium after the dominant species was positively perturbed for 500 time points. In contrast, only 39 percent of species in each network that had low trait variation had a density higher than 0.5. In all these figures, to start with, all species in a network had an initial density of *N_i_ <* 0.005, and perturbation/forcing strength of 0.5 was applied to the species with the highest degree for a duration of 500 time points. For high trait variation *σ_i_* for all species was fixed at 0.02 and for low trait variation *σ_i_* was fixed at 0.005. Initial mean trait values were sampled as given in table 1.

## 3 Discussion

In this study, we employ an approach borrowed from signal propagation in networks (18) and use eco-evolutionary dynamical modeling to evaluate the impact of network architecture and individual variation on the revival of collapsed networks. One of the main objectives of this study was to understand whether a species-specific perturbation could be transmitted across the network and could be used as a tool to revive collapsed networks in unfavorable environmental conditions.

We found that complex mutualistic ecological networks exhibited hysteresis and thus never fully recovered once environmental conditions were restored (Fig 1B-C, Fig. S1). Thus, restoring the original environmental conditions might not be a practical solution to resurrect such ecological communities back to some form of functionality (15). Therefore, understanding the dynamic behavior of these networks is critical in forecasting their future states in response to perturbation. For ecological networks that have lost functionality i.e., networks that have lost more than 90 percent of their species and/or networks that had species density ranging below a density of 0.005, we used a species-specific perturbation as a tool to steer collapsed networks to a more functional stable state (18). We picked a species that had the highest number of interactions as the focal species to perturb. In a collapsed state, an ecological network has lost its inherent capability to steer itself to some functional form under unfavorable conditions. Thus, outside intervention in some form of positive perturbation could be used to help revive an ecological network (34).

When the species with the highest number of interaction was forced positively, mutualistic networks could potentially recover even under unfavorable environmental conditions. This result was dependent on trait variation present, among other factors. We compared two different levels of trait variation. It is to be noted that the two levels of trait variation used were at the lower end of what one could expect when a network is at its undesirable low functional state (35). At higher levels of trait variation, when the most generalist species was perturbed, recovery richness after the perturbation was stopped was much higher in comparison to when individual trait variation was lower (Fig. 3B-F). However, this particular result was slightly dependent on the rate of forcing, and duration of forcing (supplementary Fig. S9). At high levels of trait variation, a large network could recover in the collapse regime i.e., *γ*_0_ *<* 1.5 (see fig. 3A-D) even at lower forcing strength (*ν <* 0.25) or duration (*T <* 200).

Trait variation in species would lead to relatively higher trait overlap when compared with low trait variation, which consequently would increase mutualistic benefits (23). As a result, as the focal generalist species is positively perturbed, this increases its density thereby increasing the overall average mutualistic strength across all the interacting species. The generalist species, thus, acts as a hub to spread the positive perturbation across most species (18), thereby reviving the network from the undesirable state to high functionality in unfavorable environmental conditions. In this particular case, individual trait variation provides the base for this spread of perturbation as well as an engine for trait dynamics to take over and aid in the resurrection, once the perturbation was stopped (Fig 3A-C, and Fig. S3-S4). At this point, one might ask why the network collapsed in the first place if a high functional state is possible even when *γ*_0_ is in the collapse regime. The reason is that the decrease in *γ*_0_ occurred too suddenly for the species to adapt and readjust the network accordingly. Perturbing a single species for a limited time buys the network time to perform this readjustment. Note also that we have assumed that each species’ trait variation remains constant even as the species declines to low density. This may be a reasonable approximation if densities are suddenly reduced. If absolute population sizes are very small, genetic drift would increase, leading to the loss of heritable trait variation, which could then speed up extinction (36). Even then, we do see that networks do recover at unfavorable conditions even at very low trait variation, *σ* = 0.005, but at different parameter ranges.

When the focal species being positively perturbed was a species with the least amount of interactions or had only a few interactions (*≤* 3), mutualistic networks did not recover, regardless of the presence of high or low variation, or network topology (supplementary Fig. S2). Thus, network resurrection was highly dependent on the type of species that was being perturbed. Ninety-one percent of the networks we analyzed (i.e., 106 out of 115 networks) had plants as the one with most interactions. In nine of the networks, it was the pollinators that had the most interactions, indicating that increasing the density of the generalist pollinator might lead to recovery. This might not be practically feasible, or sometimes this approach might not work as generalist pollinators might not be effective in pollen transfer (8). However, focusing on the generalist focal plants might be feasible as increasing the density of the focal plant could increase visitation rates of pollinators, thereby aiding the revival of entire networks over time (37, 8, 38, 39). Perturbing the species with the most interactions, or efforts to increase the restoration of the dominant species (40) could potentially “spill over” to other species. For instance, efforts to conserve and increase the numbers of endangered Golden Sun moth (*Synemon plana*), restoration of a native grassland plant *Actinidia eriantha* was necessary (41). However, focusing on a species-specific approach could have potential other problems – some closely related species might not respond the same way to disturbance as others (40, 39).

At extremely low mutualistic strength, *γ*_0_ *<* 1, large networks with low connectance hardly recover. Smaller networks with high nestedness or connectance, however, do recover even at low average mutualistic strength, *γ*_0_ *<* 1 (Fig. 4 A-C). Such a result was also dependent on the presence of trait variation. With low trait variation, the proportion of recovery richness (*N_i_ >* 0.5) was low (Fig. 4A-C), and average biomass attained was low. The end goal of such network resurrection could vary depending on adaptive management strategies, or resources available to conduct such a focused resurrection of collapsed communities (42, 43). Nonetheless, it is possible to achieve a highly functional state from a collapsed state, which is concurrently dependent on the level of degradation of the community. For instance, if *γ*_0_ *<* 1 indicates the highest level of degradation, then at low levels of *γ*_0_ *<* 1, achieving a high functional biomass state might not be possible in all scenarios (Fig. 5A-B). The knowledge of the system, the type of species interactions, and the presence of feedback loops could be beneficial in managing the recovery of such networks (42). However, purely competitive communities or communities with predation could potentially constrain the recovery of collapsed networks. For instance, changes in trophic interactions that entail the removal of predators or prey might be important for a successful recovery of vegetation (44, 45).

In translating ecological theory to restoration practices in lakes or grasslands, predictive models have included threshold theory in community dynamics (46). This could help in understanding or identifying thresholds (47), which could aid in the recovery trajectory to a desired system (15). Ecological systems without any sign of hysteresis could be recovered to a desired state when reversing the original environmental driver that caused the degradation. In the presence of hysteresis, the pathways to collapse and recovery differ markedly and have been demonstrated in a wide variety of ecological systems (16, 48, 49). In such cases, restoring the environmental conditions may not result in a community recovery and an alternative route to reviving ecological communities would be required. In such systems that are governed by feedback loops, such as the one in this study i.e., mutualistic networks, our results indicate focusing on a generalist species through positive perturbation could be beneficial to steer a collapsed network to its original functionality, even at parameter spaces that lead a functional network to collapse, i.e., *γ*_0_ *<* 1.5. This idea follows from the fact that such complex communities consist of species that are intertwined in a web of interaction networks, where propagation of perturbation could occur once a particular species is perturbed. Further research on other types of species interaction such as predator-prey systems, could provide theoretical background on understanding community resurrection from a collapse state.

## 4 Methods and Models

### 4.1 Modeling framework

We employ an eco-evolutionary modelling framework combined with concepts from signal propagation theory to understand how structural ecological network properties and eco-evolutionary factors such as heritable trait variation could aid in the resurrection of collapsed networks. We build this framework with mutualistic plant-pollinator networks because mutualistic networks have positive feedback loops that result in the emergence of characteristic alternative stable states (50, 51, 52). Signal propagation patterns in a network and their relationship with network topology have been studied in terms of how the system responds to a local perturbation on a node of a network (18). In this particular case, a local component of the network is regularly perturbed by a signal by increasing its activity, and the penetration of the signal to other parts of the network is studied to characterize the network’s overall behavior in the dynamical regime (18). We employ a similar methodology and study the interaction of network topology, evolutionary dynamics, and trait variation in the ecological systems’ components to understand the ability of ecological networks to recover from an undesirable collapsed state.

Firstly, we describe the system of eco-evolutionary equations that characterize the dynamical behavior of mutualistic plant-pollinator networks (shown here for the plant species *i* in a network, where superscript P denotes plants)(23):

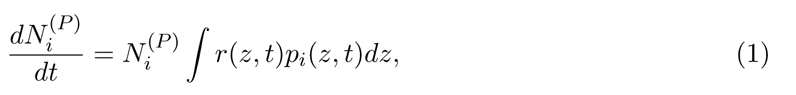

and,

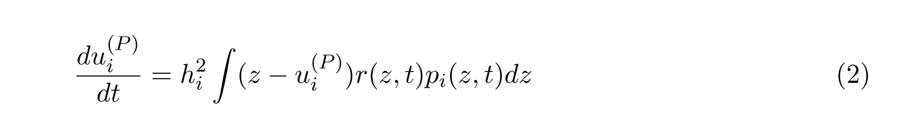

where,

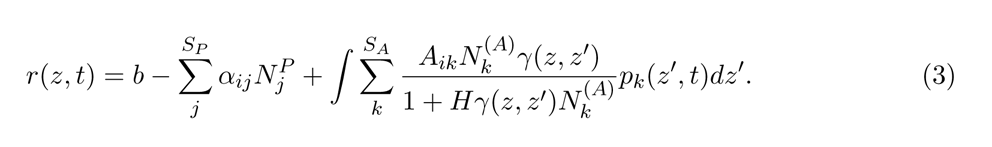

Here, equations 1 and 2 describe the dynamics of plant species density *N* ^(^*^P^* ^)^ and phenotypic trait dynamics of species *i*. *u*^(^*^P^* ^)^*i* is the mean phenotype of species *i*, and *p_k_*(*z^′^, t*) is the phenotypic distribution of animal species *k* at time *t*, where *z^′^* is the phenotype of an animal individual (for details of the derivation to the system of equations please refer to supplementary 1; the system of equations for the animal group of the network are similar as the above two but see supplementary 1). *b* is the intrinsic growth rate which was fixed at 0 for all species, indicating that species in mutualistic networks relied solely on mutualistic benefits for their survival and growth (table 1 for details). *A_ik_* is the adjacency matrix of 0 or 1 where 0 would mean no interaction between animal species *i* and plant species *k* and 1 would mean an existing interaction, *S_A_, S_P_* are the total numbers of animal and plant species in the network; *α_ij_* is the competition coefficient and the effect of competition from species *j* on species *i*; *H* is the handling time and *H >* 0, which means increases in growth rate due to mutualistic plant-pollinator interaction follows a type 2 functional response curve. This indicates that increases in mutualistic benefits of a species *i* are not linear but saturate as the interacting species density increases. *γ*(*z, z^′^*) is the Gaussian mutualistic interaction kernel given as

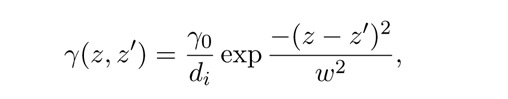

where *γ*_0_ represents the average interaction strength, and *w* designates the width of the interaction kernel, and *d_i_* is the degree of species *i*. Here, average interaction strength trades off with the species degree ensuring that species with many interactions do not become overly abundant (21, 17, 23). Finally, *h*^2^ captures the broad-sense heritable variation of the phenotypic trait of species *i*. Here, the mean phenotypic trait *u_i_* changes over time in response to changes in mutualistic interactions in the network. Species in mutualistic networks could interact via phenotypes such as the proboscis of pollinators and corolla lengths in plant-pollinator systems (30, 31). Species with similar phenotypes will gain strong mutualistic benefits in terms of growth as according to the trait-matching model (28). The above system of equations for plant-pollinator mutualistic dynamics exhibits contrasting alternative stable states w.r.t to average mutualistic strength, *γ*_0_. At a certain *γ*_0_, mutualistic networks shift abruptly from a high biomass state to an alternative low biomass state (51, 52, 23). The threshold *γ*_0_ at which such a shift occurs is also dependent on the presence of trait variation as previously shown in (23). This *γ*_0_ is central to the mutualistic interaction networks and changes in *γ*_0_ could be indirectly linked to changes in environmental conditions such as temperature (53). Changes in temperature could shift average interaction strength to a point where such a critical transition occurs. Reversing such environmental changes could be practically impossible. Even if it were possible, the system of equations could potentially exhibit hysteresis. Thus, we employ a perturbation scenario where we force one single species from a network for a certain duration of time and track the eco-evolutionary dynamics of entire networks.

### 4.2 Collapse and hysteresis regime of mutualistic networks

In our modeling framework, we assumed competition to be generally weaker than mutualism. In addition, intraspecific competition was fixed at 1 and much stronger than interspecific competition (i.e., *α_ii_ >> α_ij_*) which was drawn from a uniform distribution of *U* [0.0001, 0.001] (23, 51). Next, we evaluated the collapse regime of plant-pollinator networks by decreasing the average mutualistic strength *γ*_0_ instantaneously to a lower value. Across different model runs, we decreased *γ*_0_ sequentially from 5 to 0 in steps of 0.15. For each value of *γ*_0_, we simulate the dynamics until time 10^4^. Initial species density was fixed at 1 for all species in all the networks. We then estimated plant and pollinator total density from the last 1000 time points. The collapse threshold for species was set to 0.05, i.e., species below this density were considered to be collapsed. As the average strength of mutualistic interactions, *γ*_0_ decreased, species collapsed until the entire mutualistic networks transitioned to an undesirable collapse state. Next, we roughly estimated the collapse regime as the parameter space of *γ*_0_ where species richness remained below 90 percent of maximum possible richness for all networks. We thus then focused our next part of the analysis around this *γ*_0_ regime.

In order to estimate the hysteresis regime, we gradually increased the average mutualistic strength *γ*_0_ sequentially from 0 to 5 in steps of 0.15. For each value of *γ*_0_, we simulated ecoevolutionary dynamics until 10^4^. In contrast to the simulations of the collapse regime, here the initial starting density of each species in all the networks was ensured to be below *N* ^(^*^A,P^* ^)^ *<* 0.005 i.e., to start with species were either locally extinct or had very low density. Intraspecific trait variance was kept at moderate levels for all species, i.e., at *σ_i_* = 0.009. As *γ*_0_ increases, if a mutualistic network exhibits hysteresis, the path of recovery from a collapsed state to a fully functional state would be different i.e., networks could remain in the collapsed state even at high average mutualistic strength, *γ*_0_.

### 4.3 Perturbation regime in mutualistic networks

We perturb a single species in a mutualistic network for a certain duration. The rest of the species in the mutualistic network remain unperturbed. We chose the species to be perturbed as the one with the highest degree, i.e., the highest number of species interactions. Before we perturbed a species in a network for a certain duration when the network has transitioned to the alternative collapse state, we first used the system of eco-evolutionary equations to simulate dynamics till a network reached a positive quasi-equilibrium state. To start with, the initial species densities of all species in a network were fixed at 1, and average phenotypic species values, *u_i_*, were randomly sampled from a uniform distribution of *U* [*−*0.5, 025]. We then simulate the dynamics of the network over a time period of 10^4^ time points for an average mutualistic strength *γ*_0_ = 4. Note that this *γ*_0_ value of 4 does not fall in the collapse regime (Fig. 1). At the final time point, we take the final mean trait values of species for final simulations of network resurrection. This was done to initiate conditions that demonstrate the mutualistic network to be at a state of some kind of quasi-positive eco-evolutionary equilibrium, meaning that species densities are at the highest, and mutualist partners have adapted to each other dictated by the structure of the network. From our analysis of the collapse regime from 1, we observed that almost all networks transitioned from a high biomass state to the collapse state in the range of *γ*_0_ = [0.5, 1.5]. Knowing this, we focused our subsequent analysis of network revival around this range of *γ*_0_ values. Around these *γ*_0_ values, hysteresis existed, and thus restoring the original environmental conditions, which practically sometimes could be infeasible, did not result in the recovery of mutualistic networks (Fig. 1B). We thus evaluated whether forcing a single species in mutualistic plant-pollinator networks could instead be used to resurrect such collapsed networks. To do that, as a naive approach, we chose the species with the highest degree (be it a plant or pollinator) in each of the hundred and fifteen empirical plant-pollinator networks we analyzed. To contrast and compare, we also chose to perturb a species that instead was randomly chosen from the network that had *d_i_ ≤* 3 (see Fig. S2). We set the initial densities of all species in mutualistic networks to *N* ^(^*^A,P^* ^)^ *<* 0.005, i.e., the initial *N_i_*’s were sampled from a random uniform distribution with a range of *U* [0, 0.005]. The initial mean phenotypic values for a network were taken from the quasi-positive equilibrium when average mutualistic strength was *γ*_0_ = 4. Thus, to evaluate the temporal propagation of a perturbation and resurrection, we introduce a condition on the focal species to be perturbed with the highest degree as,

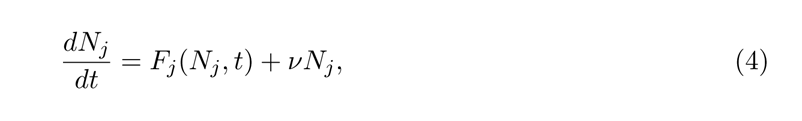

where, *F_j_*(*N_j_, t*) is right hand side term of equation 1 for species *j* (could be either plants or animals), and *ν* is the forcing strength applied for a duration of time *T*. Thus species *j* was positively perturbed for a certain duration, and at *t* = *T*, *ν* was then set to zero. In our simulations, *ν* can range from 0 to 1. This perturbation is akin to increasing the survival or fertility of a species by a certain rate, since we fix *b*, the intrinsic growth rate of species to be zero. So any *ν* value greater than 0 would essentially mean a perturbation that increases the growth of a species. We also test another specific perturbation that mimics a scenario where instead of increasing the rate of increase of a species, we continuously add some density (*ν_c_*) of the species for a duration of *T* time point. In that case (see supplementary for results), equation 4 would read as:

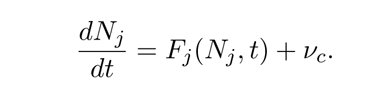

While species *j* was positively forced, the remaining (*S −* 1) species remained unperturbed (18). After the species *j* is perturbed, we then track the eco-evolutionary dynamics of all the species in the network, and evaluate for two moderate levels of trait standard deviation, *σ_i_* (high and low), whether after the duration of perturbation *T*, network resurrection took place. Since the parameter regime of *γ*_0_ where the collapse of mutualistic networks occurred ranged from as low as 0.5 to as high as 1.5, we restricted our perturbation simulations to this parameter range. In other words, for each level of *γ*_0_ that ranged from 0.5 to 1.5, and for two levels of trait variation (high and low), and for different forcing strength, *ν*, we evaluated whether forcing one single species in the collapse regime (i.e., unfavorable environmental conditions) could resurrect networks to a moderate or high functioning state. We count a species that was not directly perturbed as recovered if the density of the species reached 0.5 from a density of less than 0.005, after the perturbation was stopped. In that way, we track the proportion of species that recovered for each of the 115 plant-pollinator networks and the mean and total biomass of networks reached. For two levels of trait standard variation, high *σ_i_*was fixed for all species at 0.02, and low *σ_i_* which was fixed at 0.005. Previous studies have suggested that individual trait variation could impact the occurrence and timing of network collapses, especially in mutualistic networks. Here, we evaluated whether moderate levels of phenotypic trait variance in species mean phenotypes could impact the resurrection of collapsed networks. Note that the high level of variance in trait values sampled in this study is at the low end of the spectrum as observed in field studies of plants (54) and pollinators (55, 56). Please refer to Table 1 for details of parameter values.

In addition, for each of the empirical plant-pollinator networks, we estimated common network topological properties such as connectance and nestedness. Connectance was calculated as the total number of interactions divided by the square of the total number of species. For nestedness, we used the commonly used metric known as NODF (57). We thus then linked how network architecture could impact the process of resurrection of collapsed networks. Specifically, we asked: Do architectural constraints imposed by how species interact impact the propagation of a perturbation and thereby affect the revival of networks? Our main objective was to evaluate the role of individual trait variation, and network topology in promoting the resurrection of networks. Additionally, we also do perturbation on species intrinsic growth rate, *b*. In this particular scenario, we again focused on the species with the highest degree. *b* for all species was fixed at a value of 0. When the focal species with the highest degree was forced, we evaluated how increasing the rate of growth of the focal species could mediate in the resurrection of collapsed networks. While doing that, the rest of the S-1 species intrinsic growth rate was fixed at 0, and only the rate of growth *b* of the focal species with the highest degree was increased. While this was done, the original perturbation of *ν* continued for the duration of time point *T*. We thus evaluated whether collapsed networks could be recovered for the parameter range of *b* of the focal species with the highest degree from 0.05 to 0.5, for a range of forcing strength *ν* from 0 to 1 in steps of 0.1, for a forcing duration of *T* = 500 time steps. Individual trait variance was fixed for all species to be low at *σ_i_* = 0.005. We show this for three different types of networks with contrasting network topologies (see Fig. S6).

## Acknowledgments

GB would like to acknowledge DFG Walter Benjamin grant no BA 7974/1-1 for funding the research.

## Author contributions

GB formulated the study, ran the simulations, and analysed the model, with input from MW. GB wrote the first draft and MW contributed to the writing.

## References

[1] Scheffer, M., Hosper, S. H., Meijer, M.-L., Moss, B. & Jeppesen, E. Alternative equilibria in shallow lakes. Trends in Ecology & Evolution 8, 275–279 (1993). URL https://www.sciencedirect.com/science/article/pii/016953479390254M.

[2] Dakos, V. et al. Ecosystem tipping points in an evolving world. bioRxiv 447227–447227 (2018). URL https://www.biorxiv.org/content/early/2018/10/24/447227.

[3] Scheffer, M. *Critical transitions in nature and society* (Princeton University Press, 2009). URL https://press.princeton.edu/titles/8950.html.

[4] Pinsky, M. L., Jensen, O. P., Ricard, D. & Palumbi, S. R. Unexpected patterns of fisheries collapse in the world’s oceans. Proceedings of the National Academy of Sciences 108, 8317–8322 (2011). URL https://www.pnas.org/content/108/20/8317.

[5] Kéfi, S. et al. Spatial vegetation patterns and imminent desertification in Mediterranean arid ecosystems. Nature 449, 213–217 (2007).

[6] Kaiser-Bunbury, C. N., Vázquez, D. P., Stang, M. & Ghazoul, J. Determinants of the microstructure of plant–pollinator networks. Ecology 95, 3314–3324 (2014). URL 10.1890/14-0024.1.

[7] Kaiser-Bunbury, C. N., Muff, S., Memmott, J., Müller, C. B. & Caflisch, A. The robustness of pollination networks to the loss of species and interactions: a quantitative approach incorporating pollinator behaviour. Ecology Letters 13, 442–452 (2010). URL 10.1111/j.1461-0248.2009.01437.x.

[8] Menz, M. H. M. et al. Reconnecting plants and pollinators: challenges in the restoration of pollination mutualisms. Trends in Plant Science 16, 4–12 (2011). URL https://www.sciencedirect.com/science/article/pii/S1360138510001962.

[9] Dakos, V. et al. Slowing down as an early warning signal for abrupt climate change. Proceedings of the National Academy of Sciences of the United States of America 105, 14308–12 (2008). URL http://www.pubmedcentral.nih.gov/articlerender.fcgi?artid=2567225&tool=pmcentrez&rendertype=abstract.

[10] Scheffer, M. et al. Anticipating Critical Transitions. Science 338, 344–348 (2012).

[11] Baruah, G. Transitions and its indicators in mutualistic meta-networks: effects of network topology, size of metacommunities and species dispersal. Evolutionary Ecology 37, 691–708 (2023). URL 10.1007/s10682-023-10239-3.

[12] Baruah, G., Clements, C. F., Guillaume, F. & Ozgul, A. When Do Shifts in Trait Dynamics Precede Population Declines? The American Naturalist 193, 633–644 (2019). URL 10.1086/702849.

[13] Boerlijst, M. C., Oudman, T. & de Roos, A. M. Catastrophic Collapse Can Occur without Early Warning: Examples of Silent Catastrophes in Structured Ecological Models. PLoS ONE 8 (2013).

[14] Arkilanian, A. A., Clements, C. F., Ozgul, A. & Baruah, G. Effect of time series length and resolution on abundance-and trait-based early warning signals of population declines. *Ecology* 101, e03040 (2020). URL 10.1002/ecy.3040.

[15] Suding, K. N. & Hobbs, R. J. Threshold models in restoration and conservation: a developing framework. Trends in Ecology & Evolution 24, 271–279 (2009). URL https://www.sciencedirect.com/science/article/pii/S0169534709000470.

[16] Isbell, F., Tilman, D., Polasky, S., Binder, S. & Hawthorne, P. Low biodiversity state persists two decades after cessation of nutrient enrichment. Ecology Letters 16, 454–460 (2013). URL 10.1111/ele.12066.

[17] Lever, J. J., van Nes, E. H., Scheffer, M. & Bascompte, J. The sudden collapse of pollinator communities. Ecology Letters (2014).

[18] Hens, C., Harush, U., Haber, S., Cohen, R. & Barzel, B. Spatiotemporal signal propagation in complex networks. Nature Physics 15, 403–412 (2019). URL https://www.nature.com/articles/s41567-018-0409-0.

[19] Dunne, J. A. & Williams, R. J. Cascading extinctions and community collapse in model food webs. Philosophical Transactions of the Royal Society B: Biological Sciences 364, 1711–1723 (2009). URL 10.1098/rstb.2008.0219.

[20] Bascompte, J. & Jordano, P. Plant-Animal Mutualistic Networks: The Architecture of Biodiversity. *Annual Review of Ecology*, Evolution, and Systematics 38, 567–593 (2007). URL 10.1146/annurev.ecolsys.38.091206.095818.

[21] Bastolla, U. et al. The architecture of mutualistic networks minimizes competition and increases biodiversity. Nature 458, 1018–1020 (2009). URL https://www.nature.com/articles/nature07950.

[22] Valdovinos, F. S. Mutualistic networks: moving closer to a predictive theory. Ecology Letters 22, 1517–1534 (2019). URL 10.1111/ele.13279.

[23] Baruah, G. The impact of individual variation on abrupt collapses in mutualistic networks. Ecology Letters 25, 26–37 (2022). URL 10.1111/ele.13895.

[24] Bascompte, J., Jordano, P., Melián, C. J. & Olesen, J. M. The nested assembly of plant–animal mutualistic networks. Proceedings of the National Academy of Sciences 100, 9383–9387 (2003). URL https://www.pnas.org/content/100/16/9383.

[25] Balvanera, P. et al. Quantifying the evidence for biodiversity effects on ecosystem functioning and services. Ecology Letters 9, 1146–1156 (2006).

[26] Cardillo, M. et al. Multiple Causes of High Extinction Risk in Large Mammal Species. Science 309, 1239–1241 (2005). URL http://www.sciencemag.org/content/309/5738/1239%5Cnhttp://www.ncbi.nlm.nih.gov/pubmed/16037416: //http://www.sciencemag.org/content/309/5738/1239.full www.sciencemag.org/content/309/5738/1239.full http://www.sciencemag.org/content/309/5738/1239.full.pdf.

[27] Cahill, A. E. et al. How does climate change cause extinction? Proceedings of the Royal Society B: Biological Sciences 280, 20121890–20121890 (2012). URL 10.1098/rspb.2012.1890.

[28] Nuismer, S. L., Jordano, P. & Bascompte, J. Coevolution and the Architecture of Mutualistic Networks. Evolution 67, 338–354 (2013). URL https://onlinelibrary.wiley.com/doi/abs/10.1111/j.1558-5646.2012.01801.x.

[29] McPeek, M. A. Evolutionary community ecology. (2017).

[30] Agosta, S. J. & Janzen, D. H. Body size distributions of large Costa Rican dry forest moths and the underlying relationship between plant and pollinator morphology. Oikos 108, 183– 193 (2005). URL 10.1111/j.0030-1299.2005.13504.x.

[31] Santamaría, L. & Rodríguez-Gironés, M. A. Linkage Rules for Plant–Pollinator Networks: Trait Complementarity or Exploitation Barriers? PLOS Biology 5, e31 (2007). URL 10.1371/journal.pbio.0050031.

[32] Guimarães, P. R. et al. Interaction Intimacy Affects Structure and Coevolutionary Dynamics in Mutualistic Networks. Current Biology 17, 1797–1803 (2007). URL https://www.sciencedirect.com/science/article/pii/S0960982207020635.

[33] Chaparro Pedraza, P. C., Matthews, B., de Meester, L. & Dakos, V. Adaptive Evolution Can Both Prevent Ecosystem Collapse and Delay Ecosystem Recovery. *The American Naturalist* 198, E185–E197 (2021). URL 10.1086/716929. Publisher: The University of Chicago Press.

[34] Sanhedrai, H., et al. Reviving a failed network through microscopic interventions. *Nature Physics* 18, 338–349 (2022). URL https://www.nature.com/articles/s41567-021-01474-y. Number: 3 Publisher: Nature Publishing Group.

[35] Burkle, L. A., Glenny, W. R. & Runyon, J. B. Intraspecific and interspecific variation in floral volatiles over time. Plant Ecology 221, 529–544 (2020). URL 10.1007/s11258-020-01032-1.

[36] Nabutanyi, P. & Wittmann, M. J. Models for Eco-Evolutionary Extinction Vortices under Balancing Selection. *The American Naturalist* 197, 336–350 (2021). URL 10.1086/712805. Publisher: The University of Chicago Press.

[37] Kaiser-Bunbury, C. N., et al. Ecosystem restoration strengthens pollination network resilience and function. *Nature* 542, 223–227 (2017). URL https://www.nature.com/articles/nature21071. Number: 7640 Publisher: Nature Publishing Group.

[38] Devoto, M., Bailey, S., Craze, P. & Memmott, J. Understanding and planning ecological restoration of plant–pollinator networks. Ecology Letters 15, 319–328 (2012). URL 10.1111/j.1461-0248.2012.01740.x.

[39] Williams, N. M. Restoration of Nontarget Species: Bee Communities and Pollination Function in Riparian Forests. Restoration Ecology 19, 450–459 (2011). URL 10.1111/j.1526-100X.2010.00707.x.

[40] Lindenmayer, D. B. et al. The Focal-Species Approach and Landscape Restoration: a Critique. Conservation Biology 16, 338–345 (2002). URL 10.1046/j.1523-1739.2002.00450.x.

[41] O’Dwyer, C. & Attiwill, P. M. Restoration of a Native Grassland as Habitat for the Golden Sun Moth Synemon plana Walker (Lepidoptera; Castniidae) at Mount Piper, Australia. Restoration Ecology 8, 170–174 (2000). URL 10.1046/j.1526-100x.2000.80024.x.

[42] Suding, K., Spotswood, E., Chapple, D., Beller, E. & Gross, K. Ecological Dynamics and Ecological Restoration. In Palmer, M. A., Zedler, J. B. & Falk, D. A. (eds.) *Foundations of Restoration Ecology*, 27–56 (Island Press/Center for Resource Economics, Washington, DC, 2016). URL 10.5822/978-1-61091-698-1_2.

[43] Wainwright, C. E. et al. Links between community ecology theory and ecological restoration are on the rise. Journal of Applied Ecology 55, 570–581 (2018). URL 10.1111/1365-2664.12975.

[44] Tanentzap, A. J. et al. Seeing the forest for the deer: Do reductions in deer-disturbance lead to forest recovery? Biological Conservation 144, 376–382 (2011). URL https://www.sciencedirect.com/science/article/pii/S0006320710004143.

[45] Hidding, B., Tremblay, J.-P. & Côté, S. D. A large herbivore triggers alternative successional trajectories in the boreal forest. Ecology 94, 2852–2860 (2013). URL 10.1890/12-2015.1.

[46] Carpenter, S. R., Ludwig, D. & Brock, W. A. Management of eutrophication for lakes subject to potentially irreversible change. Ecological Applications (1999).

[47] Gao, Y., Zhong, B., Yue, H., Wu, B. & Cao, S. A degradation threshold for irreversible loss of soil productivity: a long-term case study in China. Journal of Applied Ecology 48, 1145– 1154 (2011). URL 10.1111/j.1365-2664.2011.02011.x.

[48] Liu, Q.-X., et al. Pattern formation at multiple spatial scales drives the resilience of mussel bed ecosystems. *Nature Communications* 5, 5234 (2014). URL https://www.nature.com/articles/ncomms6234. Number: 1 Publisher: Nature Publishing Group.

[49] Baruah, G., Ozgul, A. & Clements, C. F. Community structure determines the predictability of population collapse. Journal of Animal Ecology 91, 1880–1891 (2022). URL 10.1111/1365-2656.13769.

[50] Beisner, B., Haydon, D. & Cuddington, K. Alternative stable states in ecology. Frontiers in Ecology and the Environment 1, 376–382 (2003). URL 10.1890/1540-9295%282003%29001%5B0376%3AASSIE%5D2.0.CO%3B2.

[51] Dakos, V. & Bascompte, J. Critical slowing down as early warning for the onset of collapse in mutualistic communities. Proceedings of the National Academy of Sciences (2014).

[52] Jiang, J. et al. Predicting tipping points in mutualistic networks through dimension reduction. Proceedings of the National Academy of Sciences 115, E639–E647 (2018). URL https://www.pnas.org/content/115/4/E639.short.

[53] Encinas-Viso, F., Revilla, T. A. & Etienne, R. S. Phenology drives mutualistic network structure and diversity. Ecology Letters 15, 198–208 (2012). URL 10.1111/j.1461-0248.2011.01726.x.

[54] Opedal, H. et al. Evolvability and trait function predict phenotypic divergence of plant populations. Proceedings of the National Academy of Sciences 120, e2203228120 (2023). URL 10.1073/pnas.2203228120.

[55] Arroyo-Correa, B., Jordano, P. & Bartomeus, I. Intraspecific variation in species interactions promotes the feasibility of mutualistic assemblages. Ecology Letters 26, 448–459 (2023). URL 10.1111/ele.14163.

[56] Emer, C. & Memmott, J. Intraspecific variation of invaded pollination networks – the role of pollen-transport, pollen-transfer and different levels of biological organization. Perspectives in Ecology and Conservation (2023). URL https://www.sciencedirect.com/science/article/pii/S2530064423000226.

[57] Song, C., Rohr, R. P. & Saavedra, S. Why are some plant–pollinator networks more nested than others? Journal of Animal Ecology 86, 1417–1424 (2017). URL 10.1111/1365-2656.12749.

