## APPENDIX for "Reviving collapsed networks from a single species: the importance of trait variation and network architecture"

Gaurav Baruah[1*], Meike Wittmann[1]
[1]Faculty of Biology, Theoretical Biology, University of Bielefeld, Bielefeld, 33501, Germany

### 1. Mutualistic eco-evolutionary model

#### 1.1. Quantitative trait and Lotka-Volterra dynamics

We model the dynamics of pollinators and plants in an ecologically relevant quantitative trait $z$. Each individual belonging to a guild of pollinators or plants can be described with its trait $z$, but each of the species belonging to the guild of pollinators or plants are comprised of individuals with different trait values. Now the number of individuals within species $i$ at time $t$ for pollinators will be $P_i(t)$ and for the plants it will be $N_i(t)$, and the distribution of their traits within each each species $i$ can be given by a function $p_i(z,t)$ and by definition this function satisfies

$$\int p_i(z,t)dz = 1$$

at every time $t$; the limits of integration encompass the whole trait axis, which for simplicity we take to go between minus and plus infinity unless otherwise noted. $N_i(t)p_i(z,t)dz$ is then the population density of species $i$'s individuals with phenotype value between $z$ and $z + dz$ for the plants. Analogously, for the pollinators we can write $N_i^{(A)}(t)p_i(z',t)dz'$ which is the population density with phenotype value between $z'$ and $z' + dz$.

We work in the quantitative genetic limit, i.e., the trait in question is determined by many independent loci. In this case, the following important results hold (Bulmer 1980, Falconer 1981): 1) the trait distribution is normal and 2) variance of the trait does not change in response to selection,

$$p_i(z,t) = \frac{1}{\sqrt{2\pi\sigma_i^2}} \exp\frac{-(z-u_i(t))^2}{2\sigma_i^2},$$

where $u_i(t)$ is the mean trait value for the species $i$ (could be plants or pollinators) and $\sigma_i^2$ is the trait variance. In this scenario, only the mean of the trait responds to selection and the trait variance remains constant. Moreover, the distribution of the trait remains normal.

The governing dynamical equations of population dynamics can be written with Lotka-Volterra equations. The per-capita growth rate can be written as (for both plants and pollinators) shown here is the pollinator [1, 2]:

$$r(N,z,t) = b - \sum_j \alpha_{ij} N_j^{(A)} + \int \sum_k A_{ik} \frac{\gamma(z,z')N_k^{(P)}}{1 + H\gamma(z,z')N_k^{(P)}} p_k(z',t)dz' \tag{1}$$

where $b$ is the growth rate independent of competition or mutualistic benefits. $a_{ij}$ is the pairwise competition term among species belonging to each own guild. For simplicity, we are going to assume that competition is not influenced by our trait of interest $z$; $\gamma(z,z')$ is the function that captures the mutualistic interactions among individuals. Here, $z$ is the trait of an individual of a species belonging to a guild say the pollinators, and $z'$ is the trait of an individual belonging to the plants. We can take this function to be a Gaussian:

$$\gamma(z,z') = \gamma_0/d_i \exp\frac{-(z-z')^2}{w^2}$$

where, $\gamma_0$ is the average strength of mutualistic interactions and $w$ is the width that controls how strongly two individuals interact. The more similar traits of two individuals belonging to two different species, namely plants and pollinators, the stronger is the mutualistic benefit.

Equation 1 represents the per-capita growth rate of an individual with phenotype $z$ interacting facilitatively with another individual with phenotype $z'$ belonging to a species of another guild and $p_j(z',t)$ is the distribution of the trait $z'$. The integration goes over the entire trait space and summed for all the species belonging to a guild. This formulation of the model is special in the sense that growth and mutualistic interactions only depend on the phenotype $z$ but not on species identity.

Now the population dynamics of species $i$ (could be represented similarly for both plants or pollinators, but shown here for pollinators) overall trait space $z$ can be written as:

$$\frac{dN_i^{(A)}}{dt} = N_i^{(A)}(t) \int r(z) p_i(z,t) dz \tag{2}$$

Substituting equation 1 into 2 we get

$$\frac{dN_i^{(A)}}{dt} = N_i^{(A)}(t) \int \left( b - \sum_j \alpha_{ij} N_j + \int \sum_k A_{ik} \frac{\gamma(z,z') N_k^{(P)}}{1 + H\gamma(z,z') N_k^{(P)}} p_k(z',t) dz' \right) p_i(z,t) dz \tag{3}$$

We can further solve equation 3 as:

$$\frac{dN_i^{(A)}}{dt} = N_i^{(A)}(t) \left( b - \sum_j \alpha_{ij} N_j + \int \int \sum_k A_{ik} \frac{\gamma(z,z') N_k^{(P)}}{1 + H\gamma(z,z') N_k^{(P)}} p_k(z',t) p_i(z,t) dz dz' \right) \tag{4}$$

In our model, we fix intrinsic growth to be constant and independent of trait $z$.

Since the competition among species within a guild is independent of the phenotype we modelled and from $\int p_i(z,t)dz = 1$, we get,

$$\int \sum_j \alpha_{ij} N_j p_i(z,t) dz = \sum_j \alpha_{ij} N_j$$

Dynamics of the mean phenotype $u_i(t)$ of interest can then be written as and assuming in the quantitative genetic limit, shown here for the pollinators,

$$\frac{du_i^{(A)}}{dt} = h_i^2 \int (z - u_i^{(A)}) \left( b - \sum_j \alpha_{ij} N_j + \int \sum_k \frac{\gamma(z,z') N_k^{(P)}}{1 + H\gamma(z,z') N_k^{(P)}} p_k(z',t) dz' \right) p_i(z,t) dz \tag{5}$$

where $h_i^2$ is the broad sense heritability, and $\sigma_i^2$ is the genetic variance which here is equal to phenotypic variance.

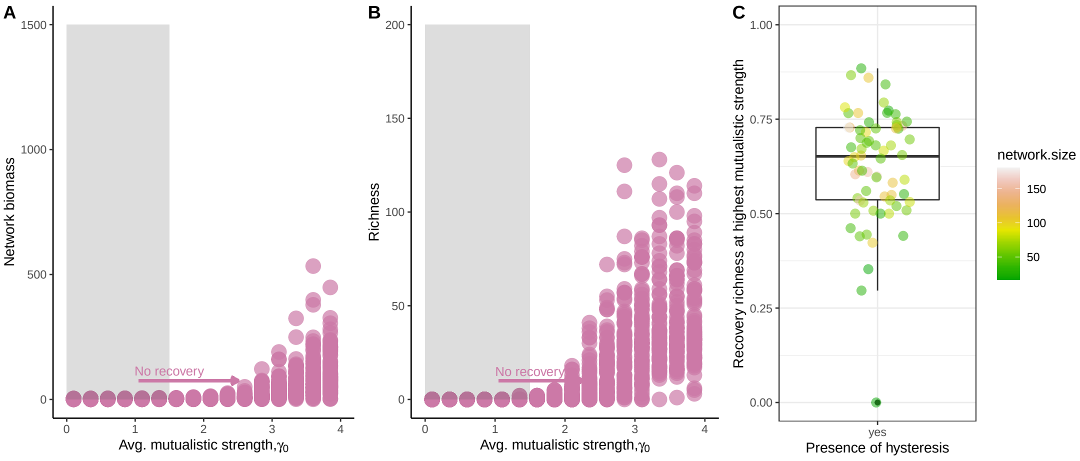


Figure S1: Hysteresis regime in mutualistic networks. (A-B) None of the 115 networks recovered to its original high functional stable state despite reverting to the original environmental conditions. (C) Recovery richness of networks at average mutualistic $\gamma_0 = 4.5$.

### References


[1] Baruah, G. The impact of individual variation on abrupt collapses in mutualistic networks. *Ecology Letters* **25**, 26–37 (2022). URL `https://onlinelibrary.wiley.com/doi/abs/10.1111/ele.13895`. _eprint: https://onlinelibrary.wiley.com/doi/pdf/10.1111/ele.13895.

[2] Barabas, G. & D'Andrea, R. The effect of intraspecific variation and heritability on community pattern and robustness. *Ecology Letters* **19**, 977–986 (2016).

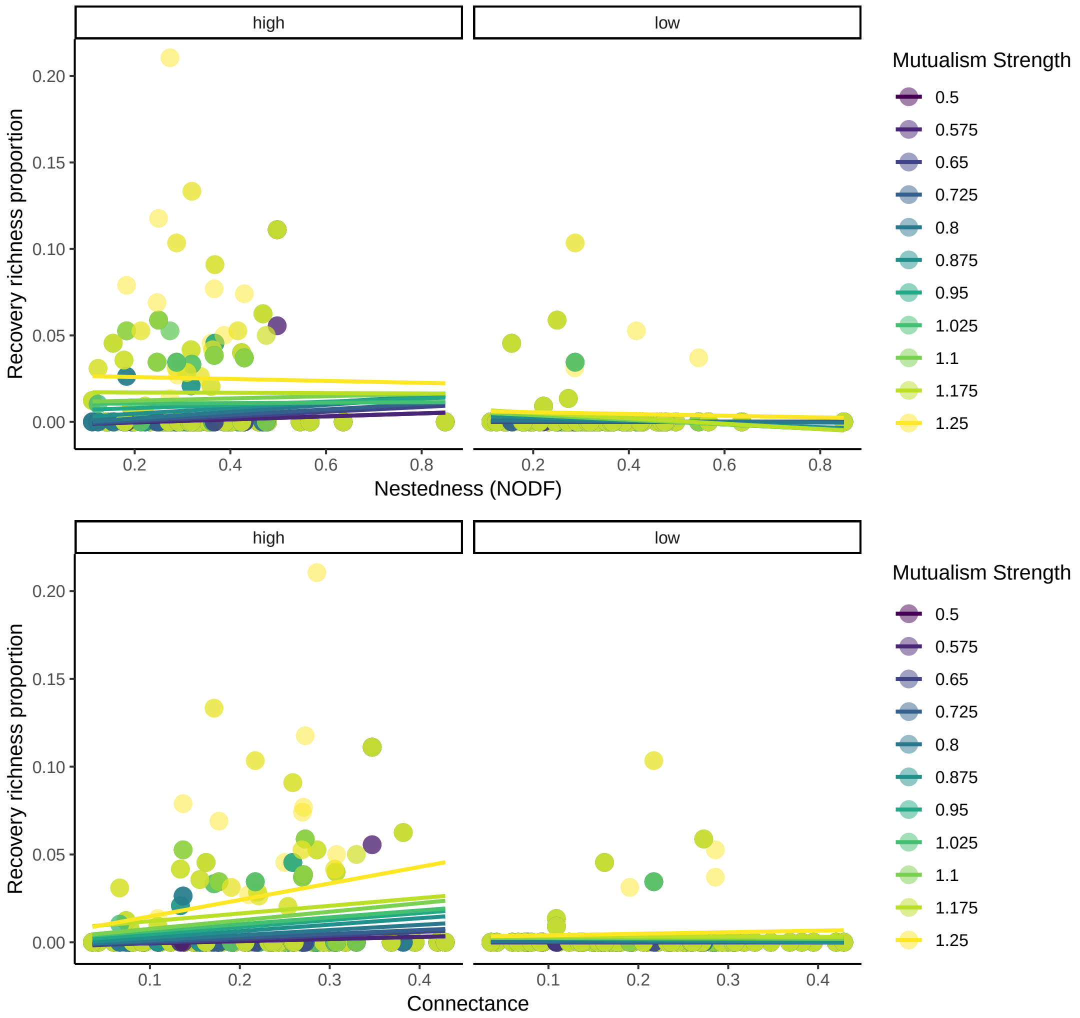


Figure S2: Recovery richness of networks as a function of Nestedness (A) and connectance (B) for a forcing strength $\nu$ of 0.5 for species that had degrees $d_i \leq 4$. Note that when a specialist species is chosen to be perturbed i.e., a species with low degree, networks did not resurrect.In the figure the faceted "high" and "low" meant high trait variation and low trait variation

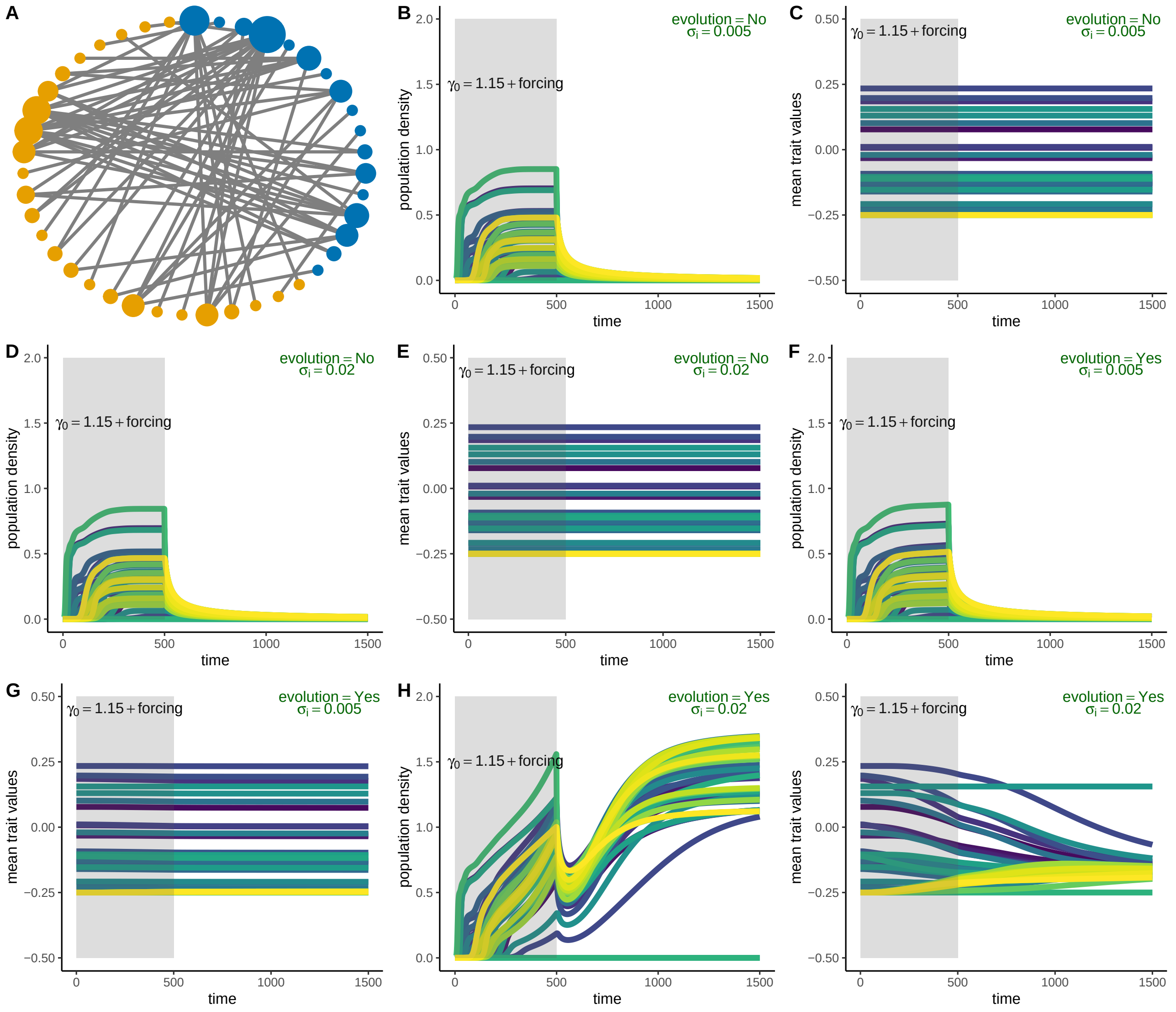


Figure S3: Impact of evolutionary dynamics on network resurrection for a mutualistic network. (A) A plant-pollinator network of 46 species. (B-C) Population dynamics and evolutionary trait dynamics of the 46 species network in the collapse regime of $\gamma_0 = 1.15$, when trait variation $\sigma_i = 0.005$ was low, and evolution was turned off, i.e., $h_i^2 = 0$ for all species. Here in B and C, at $\gamma_0 = 1.15$ only the most dominant species in terms of interactions was perturbed positively for a duration $T$ of 500 time points with a strength, $\nu = 0.5$. Network recovery fails here. (D-E) When trait variation $\sigma_i = 0.02$ was high but evolution was still turned off, i.e., $h_i^2 = 0$, network resurrection fails. (F-G) When evolution was turned on i.e., $h_i^2 = 0.4$ but trait variation was low ($\sigma_i = 0.005$), network resurrection still fails. (H-I) Only when trait variation was moderate i.e., $\sigma_i = 0.02$ and evolution turned on $h_i^2 = 0.4$, network resurrection in the collapse-regime from perturbing the most generalist species succeeds. Here in all the simulations, initial densities of all species $N_i$ were sampled from a uniform random distribution of $U[0, 0.001]$. Forcing strength $\nu = 0.5$, duration of perturbation $T = 500$.

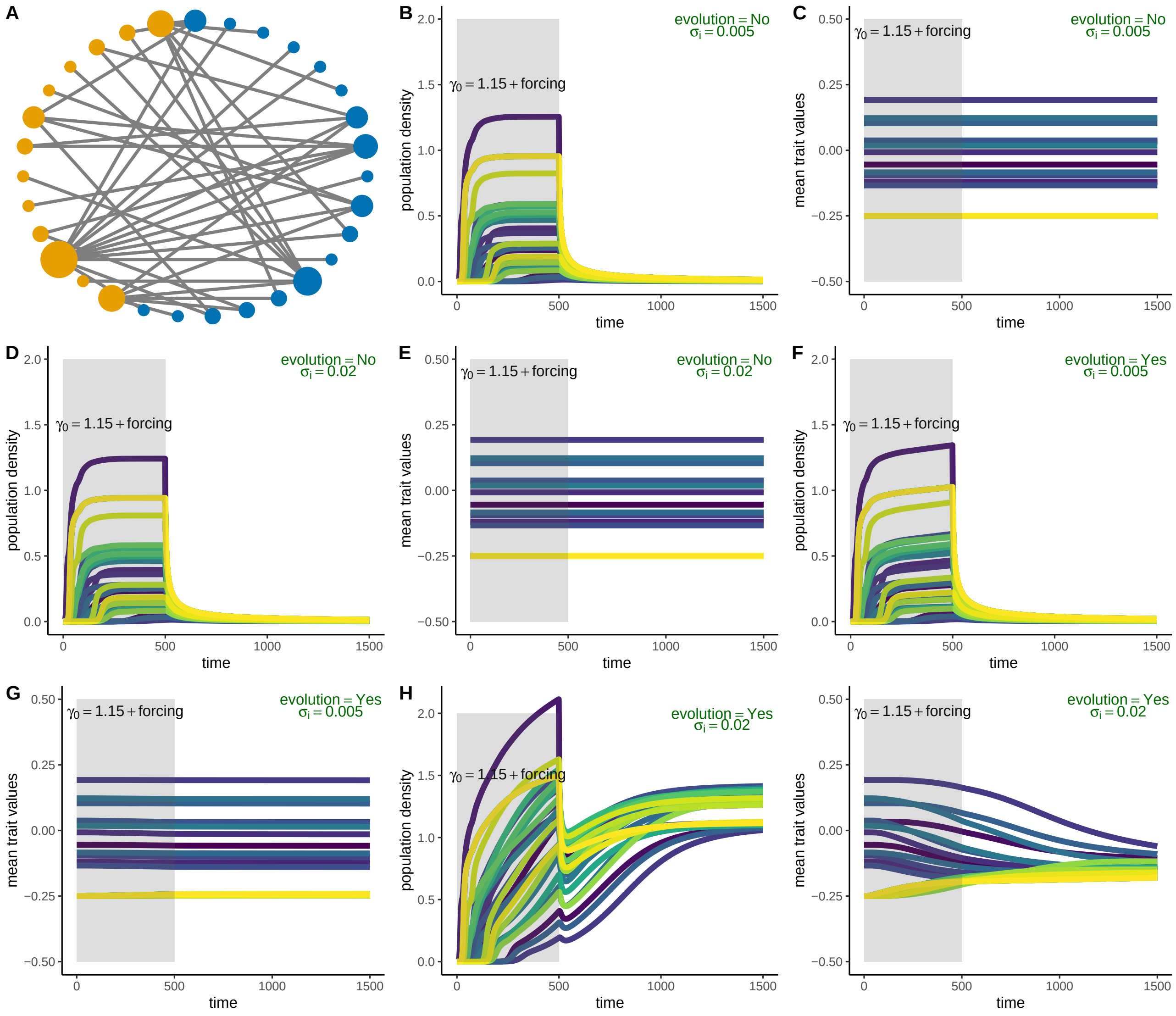


Figure S4: Impact of evolutionary dynamics on network resurrection for another example mutualistic network. (A) A plant-pollinator network of 31 species. (B-C) Population dynamics and evolutionary trait dynamics of the 31 species network in the collapse regime of $\gamma_0 = 1.15$, when trait variation $\sigma_i = 0.005$ was low, and evolution was turned off, i.e., $h_i^2 = 0$ for all species. Here in B and C, at $\gamma_0 = 1.15$ only the most dominant species in terms of interactions was perturbed positively for a duration $T$ of 500 time points with a strength, $\nu = 0.5$. Network recovery fails here. (D-E) When trait variation $\sigma_i = 0.02$ was high but evolution was still turned off, i.e., $h_i^2 = 0$. Here, network resurrection fails. (F-G) When evolution was turned on i.e., $h_i^2 = 0.4$ but trait variation was low ($\sigma_i = 0.005$), network resurrection still fails. (H-I) Only when trait variation was moderate i.e., $\sigma_i = 0.02$ and evolution turned on $h_i^2 = 0.4$, network resurrection in the collapse-regime from perturbing the most generalist species succeeds.Here in all the simulations, initial densities of all species $N_i$ were sampled from a uniform random distribution of $U[0, 0.001]$. Forcing strength $\nu = 0.5$, duration of perturbation $T = 500$.

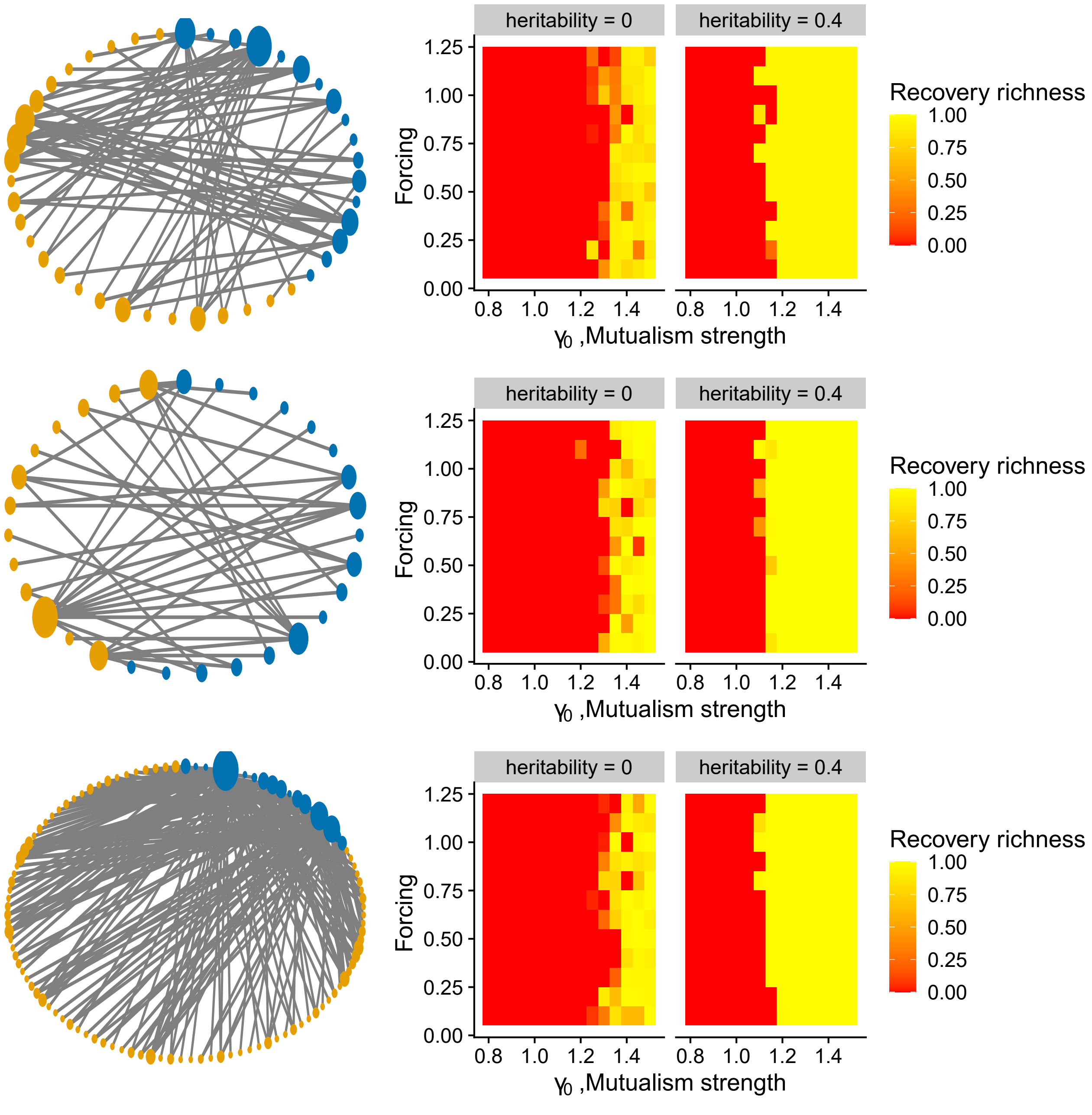


Figure S5: Comparison of network revival in the presence and absence of evolutionary dynamics on three example networks. (Top) For a network with 49 species: Network revival when forcing the species with the highest number of interactions for a wide range of forcing strength, $\nu$, and average mutualistic strength for two levels of heritability, $h_i^2 = 0$ and $h_i^2 = 0.4$. Even in the absence of heritable variation, i.e., the absence of evolutionary dynamics, network revival from a single species at unfavorable conditions could still be a possibility. (Middle) For a network with 34 species- shown for a wide range of forcing strength $\nu$ and average mutualistic strength and for two levels of heritability, $h_i^2 = 0$ and $h_i^2 = 0.4$. (Bottom) For a network with 105 species- shown here for a wide range of forcing strength, $\nu$, and average mutualistic strength and for two levels of heritability, $h_i^2 = 0$ and $h_i^2 = 0.4$. Here in all the simulations, initial densities of all species $N_i$ were sampled from a uniform random distribution of $U[0, 0.001]$. Duration of perturbation $T = 500$, and trait variation was fixed for all species at $\sigma_i = 0.02$

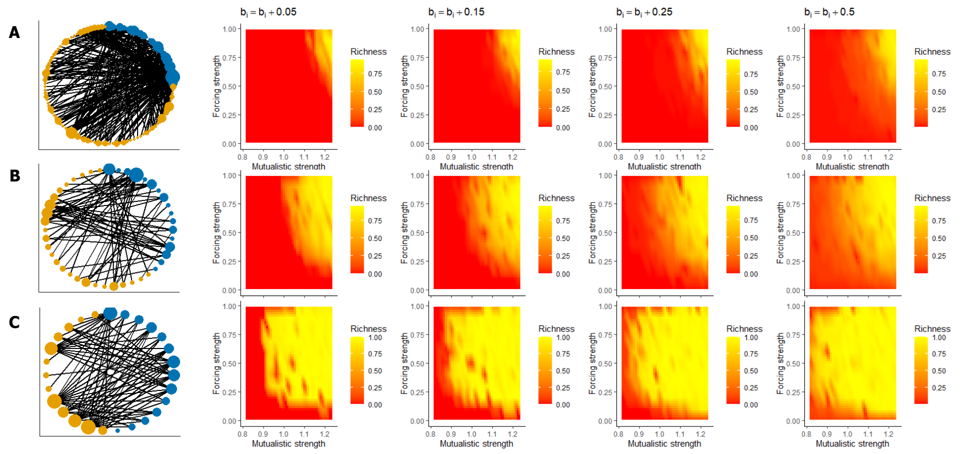


Figure S6: Parameter space of network recovery as mutualistic strength and forcing strength varies for three networks of different structures and perturbation applied to rate of intrinsic growth for the species with the highest degree. A) A network of 118 species, (B) a network of 45 species, and C) a network of 17 species. Here in (A-C), forcing was applied for a duration of 500 time points for the species with the highest degree in the collapse regime. Concurrently, species perturbation was also applied to the rate of the growth of the species being perturbed i.e., $b_i = b_i + 0.05$ means that the dominant species rate of growth was increased by 0.05, and $b_i = b_i + 0.5$ would mean that the dominant species rate of growth was increased by an amount of 0.5. Note that the species with the highest number of interactions was concurrently perturbed with a forcing strength of 0.5 as well as its rate of growth was increased.

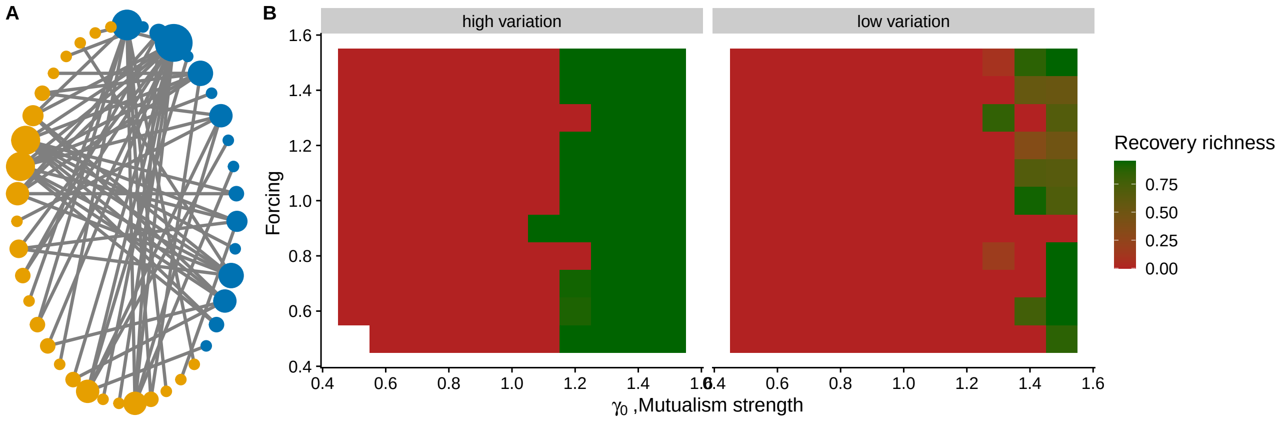


Figure S7: Parameter space of network recovery as mutualistic strength and addition of constant density of $\nu_c$ varies for a 43 species network and perturbation is applied to the species with the highest degree. Here constant addition of density of 1.2 would mean that at consecutive time points a density of 1.2 of the most generalist species is added for a duration of 500 time points. High trait variation: networks recover fully at low $\gamma_0$, whereas for low variation, networks recover only at certain $\gamma_0$ values. Initial species density was below 0.005, mean trait values were sampled from table 1 in the main-text. Low variation $\sigma_i = 0.005$, and high variation $\sigma_i = 0.02$.

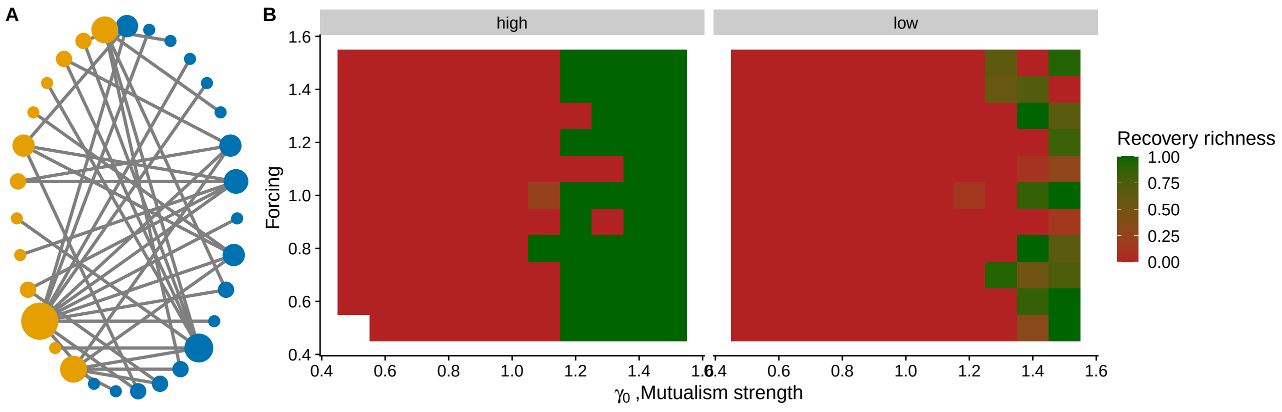


Figure S8: Parameter space of network recovery as mutualistic strength and constant density forcing strength $\nu_c$ varies for a 32 species network and perturbation applied to the species with the highest degree. Here forcing of 1.2 would mean that at consecutive time points a density of 1.2 of the most generalist species is added for a duration of 500 time points. High trait variation: networks recover fully even at low $\gamma_0$, whereas for low variation, networks recover only at certain $\gamma_0$ values. Initial species density was below 0.005, mean trait values were sampled as from table 1 in the main-text. Low variation $\sigma_i = 0.005$, and high variation $\sigma_i = 0.02$.

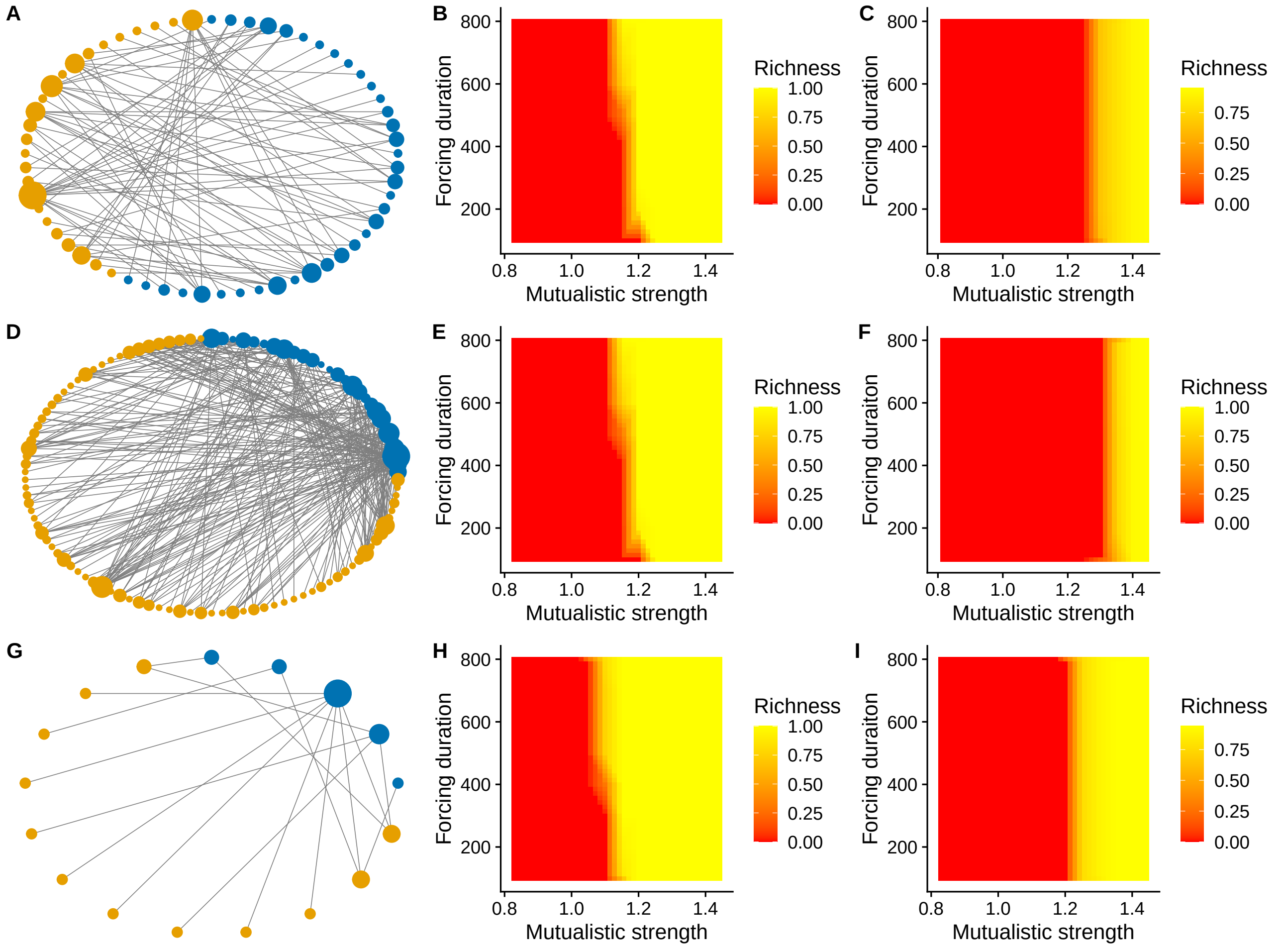


Figure S9: Parameter space of network recovery as mutualistic strength and forcing duration varies for three networks of different structures and perturbation applied to the species with the highest degree. A) A network of 61 species, connectance of 0.09, nestedness of 0.19, (B) a network of 101 species with connectance of 0.108 and nestedness of 0.221, and C) a network of 17 species with a connectance of 0.288, nestedness of 0.292. In (B-E-F) networks recover readily as forcing duration increases even at low levels of $\gamma_0$, especially when species had high trait variance in comparison to when species had low trait variance in C, F, and I. Forcing strength was fixed at 0.5, initial species density was below 0.005, mean trait values were sample as from table 1 in the main-text. Low variance $\sigma_i = 0.005$, and high variance $\sigma_i = 0.02$.
